# The Evolution of Power and the Divergence of Cooperative Norms

**DOI:** 10.1101/004416

**Authors:** Michael D. Makowsky, Paul E. Smaldino

## Abstract

We consider a model of multilevel selection and the evolution of institutions that distribute power in the form of influence in a group’s collective interactions with other groups. In the absence of direct group-level interactions, groups with the most cooperative members will outcompete less cooperative groups, while within any group the least cooperative members will be the most successful. Introducing group-level interactions, however, such as raiding or warfare, changes the selective landscape for groups. Our model suggests that as the global population becomes more integrated and the rate of intergroup conflict increases, selection increasingly favors unequally distributed power structures, where individual influence is weighted by acquired resources. The advantage to less democratic groups rests in their ability to facilitate selection for cooperative strategies – involving cooperation both among themselves and with outsiders – in order to produce the resources necessary to fuel their success in inter-group conflicts, while simultaneously selecting for leaders (and corresponding collective behavior) who are unburdened with those same prosocial norms. The coevolution of cooperative social norms and institutions of power facilitates the emergence of a leadership class of the selfish and has implications for theories of inequality, structures of governance, non-cooperative personality traits, and hierarchy. Our findings suggest an amendment to the well-known doctrine of multilevel selection that “Selfishness beats altruism within groups. Altruistic groups beat selfish groups.” In an interconnected world, altruistic groups led by selfish individuals can beat them both.

## 1. Introduction

A hallmark of human societies is that individuals are structured into coherent groups, often marked through language, dialect, or clothing (McElreath et al., 2003), and exhibit preferential treatment of in-group members (Eaton, Eswaran et al. 2011). This group structure is thought to play a large role in the extraordinary cooperation present among human societies (Hill et al., 2011; Apicella et al., 2012; Moffett 2013), and has in turn contributed to our immense success as a species (E. O. Wilson, 2012). Cooperation, however, goes hand in hand with competition. Inter-group conflict in the form of raids, warfare, and conquest are among the most important guiding forces in human cultural evolution (Tilly, 1975; Soltis et al., 1995; Turchin et al., 2013). In a world where interactions between individuals are more often positive sum (e.g., production, trade), but the most salient interactions between groups are more likely to be negative sum (e.g. warfare), the evolutionary forces selecting for individual and collective strategies are likely to push in different directions1. This raises two questions: 1) how does intergroup conflict affect individual cooperative behavior, and 2) how do collective strategies emerge from the preferences of a group’s individual members?

It is a well-documented result in the evolutionary sciences that groups that maintain cooperative or altruistic norms of behavior have an evolutionary advantage over groups with more selfish norms (Darwin 1871; Kropotkin 1905; Wilson & Wilson 2007). There has, in turn, flourished a literature regarding the mechanisms by which a group can mitigate the free-riding and parasitism that serve to undermine cooperation (Axelrod 1984; Henrich 2004; Nowak 2006; Dugatkin 1997; Santos & Pacheco 2011; Smaldino et al. 2013; Smaldino & Lubell 2014; Killingback et al. 2006; Aktipis 2004; Nowak & Sigmund 2005). This work has generally regarded selfish strategies as something to be purged or marginalized to the periphery of the group. In a world of both individual and collective interactions, however, a diversity of strategies may yield evolutionary benefits. We propose an alternative model in which individual and collective strategies can diverge, allowing cooperative norms to be maintained among the majority while a persistent minority of selfish agents yields to the group a potential evolutionary advantage. They do so via institutions that link wealth to power in collective decision making, placing the group’s collective decisions in the hands of its most selfish members. In a world where populations are increasingly interconnected, groups of individuals who can effectively cooperate with outsiders in positive sum individual-level interactions, but are collectively aggressive in negative sum group-level interactions can have a selective advantage over groups that are either uniformly cooperative or uniformly aggressive at both the individual and group levels.

Our research complements recent work indicating that cultural evolutionary models that link individual-level behavior and group-level organization can shed light on the evolution of modern social complexity (Choi & Bowles 2007; Bowles & Choi 2013; Turchin & Gavrilets 2009; Turchin et al. 2013). We present a multilevel cultural evolutionary model that illuminates the connection between prosociality, wealth, and institutions of power. The model is built on the assumption that group-level institutions (North 1990; Bowles 2006; Smaldino 2014) facilitate how strongly power in group-level decisions correlates to individual wealth, and that these institutions fall along a continuum bounded by pure democracy and pure autocracy. Individuals interact in positive sum cooperative games, groups interact in negative sum conflict games, and collective strategies are directly connected to the strategies employed by group members. Our simulations suggest that, as the global population becomes more integrated and the rate of conflict between groups increases, selection increasingly favors unequally distributed power structures, with individual influence weighted by acquired resources. These structures allow individuals to be more cooperative with outsiders through the emergence of a leadership class of the selfish. Our findings offer an amendment to the doctrine of multilevel selection, summarized by Wilson and Wilson (2007) as “Selfishness beats altruism within groups. Altruistic groups beat selfish groups.” We would add: Altruistic groups led by selfish individuals can beat them both.

### 1.1. Institutions of leadership and power

Choi and Bowles (2007) demonstrate how warfare could have facilitated the evolution of parochial altruism – i.e., the tendency to cooperate preferentially with members of the in-group while promoting hostility toward individuals of other groups. In their model, inter-group interactions resulted in war if a sufficient proportion of group members were parochial (i.e., hostile to outsiders). However, collective decisions, including the decision to go to war, are not always decided by a majority vote. In this paper, we specifically focus on the group-level institutions that translate individual proclivities for cooperation with outsiders into collective decisions related to inter-group conflict.

Humans are unique in the possession of group-level institutions that designate leaders and assign to them directive power over the group (Boehm & Flack 2010). Prior work on leadership and human evolution has focused on within-group advantages to a hierarchical power structure. Hooper et al. (2010) demonstrate that designation of a privileged individual to be responsible for monitoring and punishment can stabilize cooperation in large groups in which peer monitoring fails. In this case, we are not specifically looking at leaders as enforcers, but as individuals who make decisions on behalf of the group. A strong leadership may more efficiently consolidate resources for more effective warfare (Tilly, 1975; Turchin, 2011), and it is likely that a broad advantage of designating leaders to make collective decisions is to defray the transaction costs of decision making (Coase 1937; Williamson 1975; Williamson 1981; Dow 1987; Van Vugt et al., 2008).

This position raises the question: within whom should the group endow the power to make decisions on their behalf? Should decisions be made democratically, representing the average desires of the entire populace? Or should power in collective decisions be weighted towards a select minority? We approach this question by investigating how institutions that link power to individual resources may provide an evolutionary advantage to the group. Although institutions are often discussed in terms of promoting cooperation (Axelrod and Keohane, 1985; Bowles et al., 2003; Richerson & Henrich, 2012), institutions also can facilitate diversity in social roles (North, 1990). The power structures that emerge from institutions are group-level traits and are not reducible to properties of individuals (Smaldino, 2014). We make no assumptions about how these institutions arise or how they are maintained. We note, however, that once established, institutions may have a large degree of inertia because adopting new institutions is a vast coordination problem.

A useful simplification is to classify institutions of power along a continuum representing the degree to which power is concentrated among a subset of the group. A group whose decisions represent the average inclinations of its members can be viewed as purely *democratic.* On the opposite extreme, a group whose decisions reflect the sole inclination of a single individual can be viewed as *autocratic*, with the understanding that this operational definition ignores many of the organizational complexities associated with real autocratic regimes. A move
from democratic to more autocratic institutions of power signifies a collective decision making process that increasingly reflects the strategies of a shrinking fraction of the group.

### 1.2. Power and cooperation in an interconnected world

If today’s extant hunter-gatherer societies provide a good model for the ways in which humans lived before the rise of agriculture, then it is reasonable to assume that early humans lived according to highly democratic institutions of power. In his survey of modern-day foraging tribes, Boehm (1999) found that institutions that promoted “reverse dominance hierarchies” – in which individuals who tried to exert dominance over the group were punished – were ubiquitous. With the rise of agriculture and increased warfare, human societies became much more hierarchical, with privileged individuals deciding the fates of the communities that were increasingly organized into chiefdoms and proto-states (Carneiro, 1970; Turchin & Gavrilets, 2009). Institutions of power became less democratic and more autocratic. Turchin and colleagues (2013) have recently provided evidence for the theory that, after agriculture ended the need for nomadism and facilitated the accumulation of wealth and the rise of technology, warfare promoted the evolution of hierarchical organization of societies. Nevertheless, a puzzle remains regarding the relationship between institutions of power and cooperation. In comparison to larger-scale societies, small-scale societies are more insular, experiencing fewer interactions with outsiders. Although there is evidence that contemporary foraging societies have frequent ephemeral contact with outsiders (Fehr & Henrich, 2003), climate shifts in the Late Pleistocene – the period immediately preceding the rise of agriculture – likely led societies to be relatively isolated, both culturally and economically, in comparison to larger-scale societies (Boehm, 2012). In any case, it is uncontroversial to claim that larger-scale societies over the course of the last five millennia are characterized by substantially more contact with outsiders than are most small-scale societies, and that this is true both in terms of inter-group interactions and interpersonal interactions with extra-group individuals.

Small-scale societies are also more democratic than larger-scale hierarchical societies in terms of how collective decisions are made (at least among men – there is often dramatic inequity of power between males and females among foragers; Boehm, 1999).Yet these democratic institutions of power are associated with less cooperation among individuals who are not close kin, at least by the standards of economic games. A comparative study across 15 societies found that contributions in one-shot anonymous games of fairness (the Dictator, Ultimatum, and Third-Party Punishment Games) were correlated with the degree to which individuals tended to get their calories through market interactions as opposed to being homegrown, hunted, or fished (Henrich et al., 2010). As the authors of that study note, “Although frequent and efficient exchanges among strangers are now commonplace, studies of nonhuman primates and small-scale societies suggest that during most of our evolutionary history, transactions beyond the local group... were fraught with danger, mistrust, and exploitation” (p. 1480). In an isolated world, cooperation, particularly with outsiders, is risky business, and democratic institutions of power are widespread. In contrast, in a more interconnected world characterized by inter-group trade and war, and by less democratic institutions of power, cooperation with both in-group individuals and outsiders is comparatively more common.

In this paper we present the foundations of a theory to explain this relationship between cooperation, institutions of power, and the interconnected world. We theorize that increasing the interconnectedness of global population through interpersonal trade and inter-group conflict creates circumstances that give a selective advantage to undemocratic institutions of power, because they facilitate a power disparity that allows groups to be collectively aggressive in intergroup conflicts while the majority of individuals can be highly cooperative, increasing the overall resources available to the group. We support this theory with agent-based model simulations.

## 2. The Model

We consider a simplified world of agents, each of whom belongs to a group and is marked so as to be able to distinguish in-group members from outsiders. Model simulations run in the absence of group markers are presented in the Supplementary Information. The world initially contains *M* groups, each of which is initialized with *n* agents. Individuals pair up and have individual-level interactions which represent opportunities for cooperation or exploitation, and as modeled as a Prisoner’s Dilemma (PD) game. The relevant measure of interconnectedness for individual-level interactions is the *rate of population mixing, m,* which is the degree to which individuals are likely to have interactions with partners from outside their own group. After a round of individual-level interactions there is the potential for group-level interactions. During this, groups are randomly paired and engage in a Hawk-Dove (HD) game, which is a classic model for inter-group cooperation, exploitation, and conflict. Here the relevant measure of interconnectedness is the *rate of conflict, γ,* which is the probability that a round of group-level interactions occurs for each round of individual-level interactions. Agents die if their resources fall to zero or probabilistically with a rate that increases with age, and a group may go extinct if all of its member agents die. Otherwise, payoffs from both individual- and group-level interactions are accumulated by individuals and used for (asexual) reproduction, such that the probability of reproducing increases with acquired resources. Offspring are born into the same group as their parents, and remain there throughout their lives. We also experimented with migration as a robustness check, as it can be an important factor in evolutionary dynamics; we will present these results later. Our model structure bears some resemblance to a framework introduced by Simon and colleagues (2012). Their analyses indicate that due to the complexity of its structure, an analytical approach to our model is not feasible, and that simulations are required.

In addition to their group markers, each agent possesses two heritable traits, *p*_*in*_ and *p*_*out*_, representing the probability of cooperating in the individual-level PD game with an in-group or out-group co-player, respectively. At the beginning of each simulation, each agent is initialized with *r*_0_ resources and an age of zero. Each agent is also initialized with values for *p*_*in*_ and *p*_*out*_, each of which is a real number randomly and independently drawn from a uniform distribution in [0, 1).

### 2.1 The distribution of power as cultural institution: powergini

Each group is defined by a group-level institution relating individuals’ resources to power in collective decision making during group-level interactions. These institutions are exogenously imposed on each group and do not change. At the beginning of a round of group-level interactions, all the *n* individuals in a group *j* are ranked according to their accumulated resources, such that the agent *i* with the most resources has *rank_ij_* = 1 and the agent *h* with the least resources has *rank_hj_* = *n*. The distribution of power within a group stems from a fixed cultural institution, implemented as the parameter *power skew, σj*, which translates an agent’s ranking into the weight of its influence in collective decision making, so that the weight for an agent *i* in group *j* is given by:

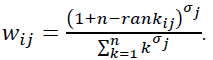

While the *power skew* is a quantified metric determining how power is distributed within a group, its scale does not have an immediately intuitive meaning in the context of real human groups. For the sake of greater explanatory power, we adopt the Gini coefficient (Damgaard & Weiner, 2000), a well-known statistic often used as a measure of inequality within a population, to characterize and index a group’s distribution of power. We adopt this metric to represent the disparity among individuals’ decision-making power within a group, and refer to this quantity as the group’s *powerGini.* To compute this statistic for each group, we assumed that the initial number of agents within a group, *n*, could be strictly ranked in terms of acquired resources (although initially they all had equal resources, this would not be case for the majority for the simulation run). We could then calculate each agent’s decision weight, *w_ij_*, as described above. The group’s *powerGini* is then given by:

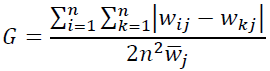

where 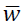 is the average decision weight in the group. A completely democratic group has *powerGini* = 0, while groups grow increasingly autocratic as *powerGini* approaches 1 (the algorithm used to ensure a nearly uniform distribution of group *powerGini* in the population is given in the Supplementary Information). Throughout the rest of the paper, the distribution of power in each group *j* is indexed as, and referenced by, its *powerGini_j_.*

### 2.2. Individual-level interactions

At the beginning of each time step, agents are paired using the algorithm described below, and each pair then plays the PD game for resource payoffs.

#### 2.2.1. Pairing algorithm

Let all the agents in a group *j* be represented by a vector **X_j_** = [*x_1j_, x_2j_*,…,x_nj_] where group *j* has *n* agents. For each group, the order of elements is randomized using a pseudorandom number generator. A new vector, **Y** = [*x_1_, x_2_*, … *x*_*N*_], is then created by concatenating each vector **X_j_** for all *M* groups in a random order. The vector **Y** therefore contains all the live agents in the simulation, clustered by group. The elements in vector **Y** are then shuffled such that each agent *i*, with probability *m*, swaps positions with a randomly selected agent in **Y**. Finally, starting with the first agent in the newly shuffled **Y**, each agent with an odd-number index is paired with the next (even-numbered) agent in the vector. When there are an odd number of total agents, one agent will not play the PD game. Thus, when *m* = 0, almost all pairings occur between agents of the same group, and when *m* = 1 pairings are completely random.

#### 2.2.2. Agent games

Each pair of agents play a standard prisoner’s dilemma (PD) game in which the payoff for mutual cooperation is the reward *R*, the payoff for mutual defection is the punishment *P*, and defecting against a cooperator results in the temptation *T* for the defector and the sucker’s payoff *S* for the cooperator. A PD game is defined when *T > R > P S* and *R* > 2 (*T* + *S*), and is a positive-sum game. The probability that an agent cooperates is determined by whether its co-player is a member of its group, as each agent *i* has two heritable traits corresponding to the probabilities of cooperating with in-group (*p*_*in*_) and out-group (*p*_*out*_) coplayers. Each agent then chooses a move probabilistically and collects payoffs, which can accumulate and be used for reproduction. Values for payoffs and other model parameters are given in Table 1.

**Table I.** Model parameters.

| Entity | subfamily | Description | Symbol | Value(s) |
| --- | --- | --- | --- | --- |
| Global | Initialization | Number of groups | $M$ | 50 |
| | | Initial number of agents per group | $N$ | 50 |
| | | Agents' initial resources | $r_0$ | [14,24]* |
| | Reproduction | Maximum reproductive rate | $K$ | 0.15 |
| | | Standard deviation of mutation | $SD_m$ | 0.02 |
| | | Reproduction growth rate | $B$ | 0.2 |
| | | Cost of reproduction | $cost\_R$ | [15,25] † |
| | | Amount of resources for maximal reproductive rate | $\rho$ | [30,50] ‡ |
| | Death | Baseline probability of death | $\delta$ | 0.001 |
| | | Max age | $maxAge$ | 50 |
| | Agent games | Population mixing rate | $m$ | [0, 1] |
| | | PD Game payoff matrix | $T, S, R, P$ | $T = 2, S = 0, R = 1.5, P = 0.5$ |
| | Group games | Rate of conflict – the probability of a round of between-group games for each round of between-agent games | $\gamma$ | [0, 1] |
| | | The returns to scale, based on total resources, for intergroup conflict (hawk-hawk) | $q$ | 1 |
| | | HD Game payoff matrix | $v, c$ | $v = 2, c = -3$ |
| | Migration | Probability of migration | $\mu$ | 0 |
| Agent |  | Group ID |  |  |
| | | Accumulated resources | $r_i$ | |
| | | Age | $ai$ | |
| | | Resources rank | $rank_i$ | |
| | | Probability of in-group cooperation | $p_{in}$ | |
| | | Probability of out-group cooperation | $p_{out}$ | |
| Group | | Power skew | $\sigma_j$ | |
| | | $powerGini$ | | |

\* $r_0 = cost\_R - 1$ † $cost\_R = 15 + (10 \cdot \gamma)$ ‡ $\rho = 2 \cdot cost\_R$ , where † and ‡ reflect the calibration of simulation parameters to the rate of conflict so that the expected reproductive value from each step of the experimented was the same across all experimental runs.

### 2.3. Group-level interactions

A round of group-level interactions occurs with probability γ, the rate of conflict. In this case, groups are paired randomly and play a HD game. If there is an odd number of extent groups, one group will not be paired. Each group *j’*s move is determined by the composition of its member agents and by its power skew, *σ_j_*, which defines the agents’ weights of influence.

We explicitly connect collective behavior to individual strategies.2 We assume that playing dove in an inter-group interaction is akin to cooperation in an individual-level interaction, and that agents’ strategies for individual-level interactions with out-group co-players are representative of their inclinations in group-level interactions. The probability that a group plays dove is related to the out-group prosociality of its individual constituents, but in more autocratic (less democratic) groups, individuals with more acquired resources have a greater influence. Specifically, the probability that a group will play dove is given by:

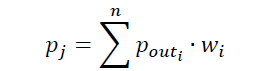

We assume that groups interact because they are both interested in some resource, *v*. If both groups are cooperative and play dove, they split the resource and each receive *v*/2. If one group plays dove and the other aggresses and plays hawk, then the aggressive group takes the entire resource and the cooperative group capitulates and gets nothing. If both groups play hawk, then there is active conflict for the resources, which is costly for the loser. The winner of the conflict receives the payoff *v*, but the loser receives a negative payoff –*c*, where *c* > *v*. Hence, active conflict is risky, and is negative-sum. Each group’s members share equally in the costs and benefits of large-scale inter-group conflicts (e.g., all agents receive *v*/2 resources when both groups play dove). This is a simplifying assumption, as individuals may differentially share costs and benefits through status or behavior (Mathew & Boyd, 2011). The winner of an active conflict is determined probabilistically. We assume that larger groups with more resources will tend to dominate smaller and poorer groups. The returns to scale are specified using methods adopted from Hirschliefer (2000). In an active conflict between two groups (labeled 1 and 2 here), the probability that group 1 will win is given by:

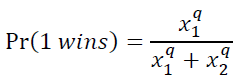

where *x*_*j*_ is the total resources of all the agents of group *j* (determined by summing the resources of all the group’s agents), and *q* represents the returns to scale. For our simulations, we assumed constant returns to scale, *q* = 1.

### 2.4. Death

An agent dies if its resources are less than or equal to zero. An agent may also die from disease or happenstance, the chance of which increases with age. Mathematically, the chance of such a death is given by:

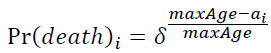

where *δ* is the baseline probability of death, *a*_*i*_ is the number of time steps for which the agent has been alive, and *maxAge* is the age at which the probability of death becomes one. For our simulations, we used *maxAge* = 50, with a average lifespan of 31.5 in the absence of resource-related deaths. Note that our agents have neither developmental nor infertile periods, so this lifespan represents active adulthood only.

### 2.5. Reproduction

An agent with sufficient resources (*r*_*i*_ > *cost_R*) reproduces probabilistically, losing *cost_R* resources in the process.3 A child agent is born into the same group as its parent, and begins with *r*_*0*_ resources.4 An agent with sufficient resources reproduces with probability given by the logistic function:

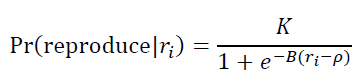

where *K* is the maximum probability of reproduction, *B* is the rate of growth, and *ρ* is the amount of resources corresponding to maximal growth in reproduction probability. In the experiments to follow, *ρ* and *cost_R* are calibrated to the rate of conflict so that the expected reproductive value from each step of the experimented was the same across all experimental runs.

An individual’s cooperative tendencies (*p*_*in*_ and *p*_*out*_) are vertically transmitted (Boyd & Richerson, 1985) from the agent’s parent with the potential for error (mutation). Mutation occurs through the addition of an error term to both *p*_*in*_ and *p*_*out*_. The error term is calculated independently for each parameter and is drawn from a Gaussian distribution with mean of zero and standard deviation *SD*_*m*_ = 0.02. Both *p*_*in*_ and *p*_*out*_ were bounded in [0, 1].

### 2.6. Migration

At each time step, *μN* agents are randomly selected, and each is reassigned to a randomly selected group. A migrating agent forsakes its former group identity and joins the new group, including associated group markers. Because destination groups are randomly selected from among extant groups (those groups with a nonzero population), a “migrating” agent could hypothetically remain in the same group. For most of the simulations presented, *μ* = 0.

## 3 Simulation Results

We ran 200 replicate simulations for each scenario reported, and the results presented are averages from across those runs. Simulations were run for 300 time steps, which was sufficient time for the distribution of agents across groups to reach a steady-state5 and for cultural, but not genetic, evolution to occur (Perrault, 2012). The values of fixed parameters and the ranges of tested parameters are summarized in Table 1.

### 3.1. Population-level results

We first consider the behavior of the average rates of in-group and out-group cooperation (i.e., the average values for *p*_*in*_ and*p*_*out*_) in the population in response to population mixing and inter-group conflict. Averaging across all rates of conflict, we obtained the result that increased population mixing selects for an increased in-group bias in individuals’ cooperative tendencies (Fig. 1a). Competition between groups, in this case in the form of conflict, creates multilevel selection pressures that reward groups who can collectively increase their total resources, which, in the absence of considering institutions of power, is best accomplished through cooperation with in-group members and defecting against out-group individuals (Hammond & Axelrod, 2006; Choi & Bowles, 2007).

**Figure 1.**
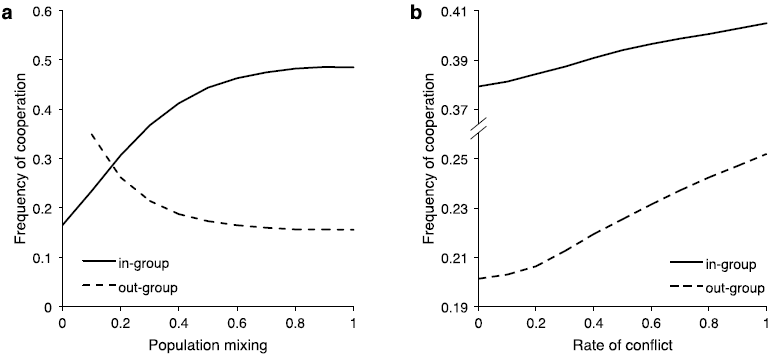
Population structure influences cooperation. The average in-group and out-group cooperation strategies as a function of (a) population mixing, averaged across rates of conflict, and (b) rate of conflict, averaged across rates of population mixing.

Averaging across all rates of population mixing and focusing on the rate of conflict indicates that increased inter-group conflict increases the rates of both in-group and out-group cooperation, and has a larger effect on out-group cooperation, effectively decreasing the average in-group bias in the population (Figure 1b). This appears to contrast with other work indicating that conflict should select for preferential cooperation with in-group individuals and increased antagonism toward outsiders (Choi & Bowles, 2007; Gneezy & Fessler, 2012). However, this prior work did not account for the presence of institutions of power. As the rate of conflict increases, the premium on generating resources as a group becomes greater. It is therefore important for group success that members maximize their productivity, even if doing so means cooperating with outsiders from groups with whom their group may eventually be in conflict. This is consistent with empirical results indicating the ascendance of cosmopolitan strategies (i.e., cooperation with outsiders) in the presence of group-level conflict (Buchan et al., 2009; Pan & Houser, 2013). As we will demonstrate, it is the presence of institutions of power makes this rise of cosmopolitanism possible.

### 3.2. Conflict influences institutional group differences

Here, we consider differential influence of increased rates of conflict on groups with differently skewed institutions of power. Figure 2 portrays three heatmaps, each corresponding to a different group-level variable. For each graph, the *x*-axis refers to the *powerGini* of a given group, varying between more democratic groups at the low end and more autocratic groups on the high end. The *y*-axis refers to the rate of inter-group conflict. A useful interpretation is that each row, corresponding to a given rate of conflict, is a portrait of a single condition of the model, in which different group-level institutions can be compared.

**Figure 2.**
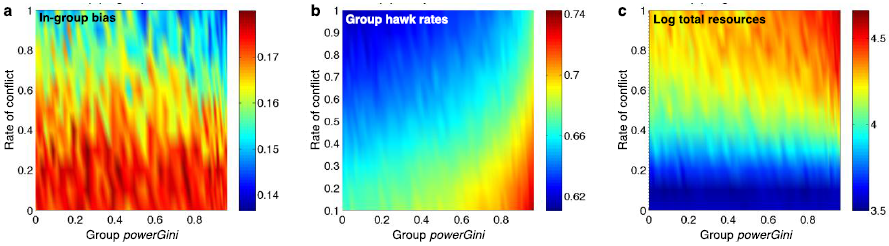
Rate of conflict differentially affects groups depending on power structure. Heatmaps indicating the average (a) in-group bias among group members, (b) rate at which groups played Hawk in inter-group interactions, and (c) the log_10_ total resources of the group, taken as a function of the rate of conflict across all groups as identified by their *powerGini*. All data is averaged across rates of population mixing.

As demonstrated above, in-group bias (defined as the average *p*_*in*_ − *p*_*out*_) decreases with the rate of conflict, as *p*_*out*_ increases faster than *p*_*in*_. The upper right corner of Fig. 2a indicates that under high conflict rates, groups with more autocratic power structures evolve less in-group-biased cooperative strategies than do those with more democratic institutions. This is because democratic groups cannot afford to be too cooperative toward outsiders in part because universal cooperative norms would translate to playing dove too often in inter-group interactions and leave them open to exploitation (in addition to the risks of exploitation in individual-level interactions), whereas more autocratic groups may be sheltered from the group-level effects of cosmopolitan norms on their collective strategies. In more autocratic groups, the individuals who are the least cooperative and therefore control the most resources (Nowak, 2006) will have more influence on group-level decisions regarding conflict. Democratic groups, on the other hand, must be more parochial, in part because being less cooperative toward out-group individuals leads to more aggressive collective behavior in inter-group interactions. This connection between individual and collective behavior is weakened in autocratic groups, allowing them to simultaneously evolve more cosmopolitan interpersonal cooperation strategies (Fig. 2a) and more aggressive collective strategies (Fig. 2b).

A more aggressive strategy means that a group will be less subject to exploitation by other groups, but also creates an increased opportunity for active conflict, which can be quite costly to the loser. More hawkish collective strategies will only benefit the group if it controls enough resources so that it has a higher than average chance of winning active conflicts. When the rate of conflict is high, there are more total resources available in the population, due to the productive and technological benefits associated with inter-group trade and warfare. More important to our argument is that when the rate of conflict is high, more autocratic groups control more total resources compared with more democratic groups (Fig. 2c). This allows these groups to benefit from being more hawkish.

### 3.3. An interconnected world selects for less democratic institutions of power

Fitness is, ultimately, measured by reproduction. When either population mixing or the rate of conflict is low, a group’s institution of power has no significant influence (statistical or otherwise) on its population. However, when both population mixing and the rate of conflict are high, the log-transformed group population sizes are positively correlated with the group *powerGini* (Figure 3).

**Figure 3.**
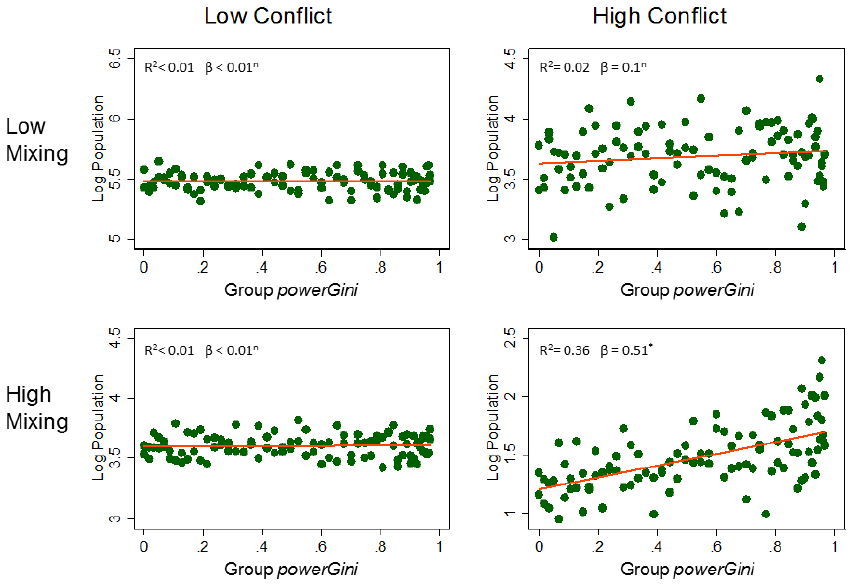
A relationship between reproductive success and undemocratic of power emerges only when both population mixing and the rate of conflict are high. Scatterplots depict the log of groups’ populations against their *powerGini* for low and high population mixing (0.1 and 1.0, respectively) and low and high rates of conflict (0.1 and 1.0, respectively). This correlation was only statistically significant when both mixing and conflict are high. Plot includes a bivariate linear model: logPopulation = β.*powerGini+*ε*.* * denotes β significant at the p<0.01 level; n = not statistically significant

To examine this phenomenon in more detail, we adopt a convenient and simplifying measure of institutional fitness, the mean population *powerGini*, which is calculated by averaging over every individual in the population and considering the *powerGini* of that individual’s group. Mathematically, this equals 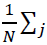, where *N* is the total number of individuals in the population and the sum is over each group *j*. Figure 4 shows that population mixing and intergroup conflict interact multiplicatively – increasing *both* mixing and conflict rates yields a selective advantage to less democratic groups, as individuals in those groups out-reproduced those in more democratic groups, but neither condition is effective in the absence of the other. Population mixing and intergroup conflicts are, as such, both necessary but not sufficient conditions to yield an evolutionary advantage to less democratic institutions of power.

**Figure 4.**
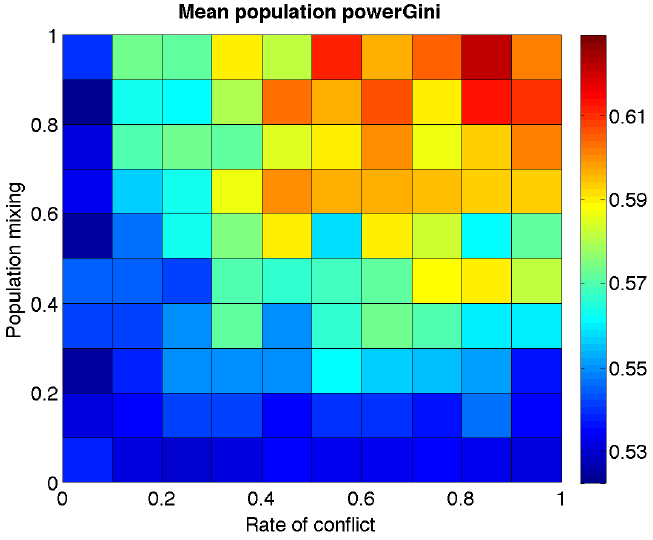
Population mixing and inter-group conflict select for undemocratic power structures. Each cell of the heat map represents a different combination of the Rate of Conflict (x-axis) and Population Mixing (y-axis) parameters. The color of the represents the mean population *powerGini* across the 200 simulations using that parameter combination. Increased population mixing and rate of conflict multiplicatively give a selective advantage to less democratic groups: individuals in those groups out-reproduced those in more democratic groups.

### 3.4. Institutions of power and the divergence of cooperative norms

Our claim is that more autocratic institutions of power succeed in a more interconnected world because they allow the less cooperative individuals to make group-level decisions regarding conflict, allowing other individuals to be more cooperative. To examine this claim in more detail, we looked at the average levels of in-and out-group cooperation found among the most and least cooperative members of each group. Figure 5 presents a series of scatterplots depicting the average 1^st^ through 99^th^ percentile distributions of the in-and out-group cooperation strategies, as a function of group *powerGini* (population-level analyses showing these percentile effects as a function of both the rate of conflict and the rate of population mixing are presented in the Supplementary Information). The in-group and out-group strategies of the least cooperative individuals in each group decline with *powerGini* (i.e., the 1^st^ and 10^th^ percentiles of cooperation are declining as groups become less democratic). At the same time, the strategies of the most cooperative individuals in each group *increase* with *powerGini* (i.e., 90^th^, and 99th percentiles of cooperation are increasing as groups become less democratic). These plots show a clear effect: Undemocratic institutions of power widen the distribution of cooperative strategies, making the least cooperative individuals more selfish, and the most cooperative individuals more cooperative.

**Fig. 5.**
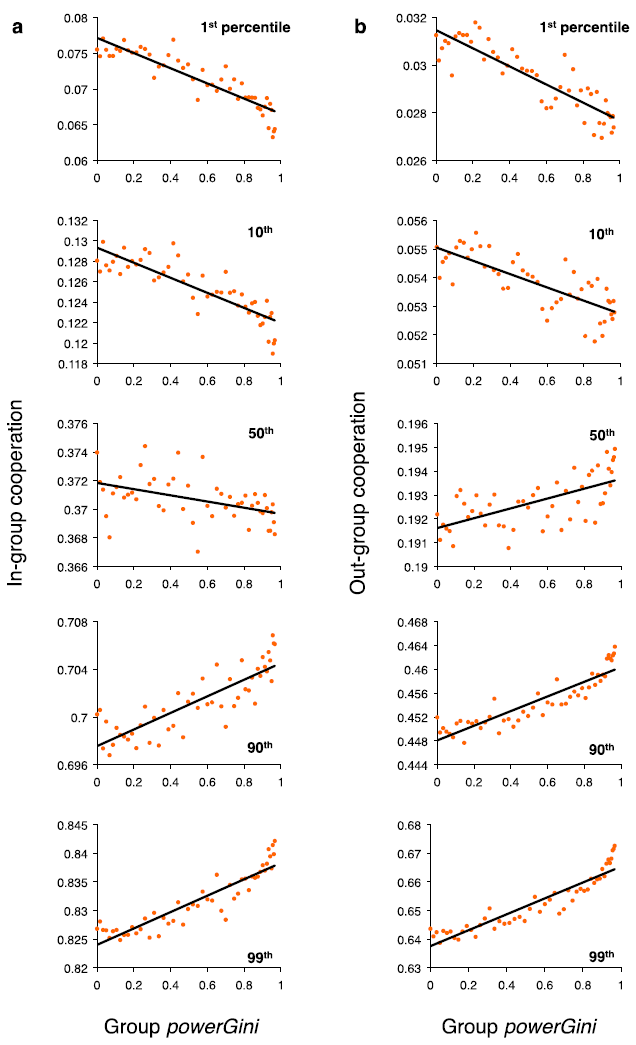
Undemocratic power structures widen the distribution of cooperation strategies, making the least cooperative individuals more selfish and the most cooperative individuals more altruistic. Scatterplots depicting the mean 1^st^ through 99^th^ percentile of the distributions of (A) in-group and (B) out-group cooperation strategies.

### 3.5. Robustness to agent movement

In many species, group-level effects on selection are negligible, because high dispersal and migration rates eliminate cohesive group structure. This is not the case for humans, however. Indeed, there are numerous psychological and cultural mechanisms in place to maintain affiliative identities among human groups (Durham 1991; Chudek & Henrich 2011), and these identities act as powerful psychological motivators for social behavior (Tajfel et al. 1971; Brewer 2004). Nevertheless, humans do sometimes change their group identity through conversion or migration, and this destabilization of group cohesion can act as a powerful force in cultural evolution (Boyd & Richerson 2009; Hirschman 1970; Powell et al. 2009).

In our baseline model, individuals interacted with members of other groups, but never changed their group identity, reflecting the fact that cultural identities are strong and robust to encounters with out-group members. It is important, however, to ensure that our results are robust to a stronger type of migration, in which individuals can change group identities (including group markers) so that their fates in inter-group interactions are tied to their adopted rather than natal group. We ran simulations in which the probability of migration *μ* = 0.01. We found that while migration added a small amount of noise, the results were qualitatively unchanged.6

## 4. Discussion

As the model’s population becomes more interconnected and the opportunities for individual and collective interactions with outsiders increases, selection will increasingly favor undemocratic institutions of power. As power in our model is concentrated within a shrinking minority of a group’s wealthiest members, these institutions facilitate both an increase in the variance of cooperative traits within the group and, in turn, a divergence of collective behavior from average individual member behavior. These institutions yield groups in which the most selfish, parochial members make collective decisions on behalf of a cooperative, cosmopolitan populace. Our model demonstrates a mechanism through which a “leadership class of the selfish” emerges, in effect insulating the majority of a group’s members and allowing the evolution of more cooperative out-group strategies.

Our results impact broadly on the literature on the evolution of cooperation in humans. This literature has been generally focused on the elucidation of mechanisms that can shift the balance between cooperators and defectors to promote the evolution of cooperative strategies. These mechanisms include reputation (Nowak & Sigmund, 2005; Smaldino & Lubell, 2014), punishment (Boyd & Richerson 1992; Bowles & Gintis, 2004), unproductive costs (Aimone, Iannaccone et al. 2013), and active avoidance of non-cooperators (Aktipis, 2004; Pepper, 2007). These mechanisms are costly and as such minorities of non-cooperators typically remain, however marginalized. The peripheral membership of non-cooperators, generally viewed as parasitic, is, within the context of our model, a potential asset that is selected *for* at the group level if their drag on interpersonal production is outweighed by their contribution towards optimizing the group’s collective inter-group strategy. Consequently, we propose an addendum to the widely accepted view concerning the evolution of cooperation and altruism in structured populations, summarized by Wilson and Wilson (2007) as “Selfishness beats altruism within groups. Altruistic groups beat selfish groups.” We would add that in an interconnected world, altruistic groups led by selfish individuals can beat them both.

In the absence of group-level interactions, groups with the most cooperative members will outcompete less cooperative groups, while within any group its least cooperative members will be its most successful7 (Nowak, 2006). The advantage to less democratic groups rests in their ability to allow most of their members to cooperate – both among themselves and with outsiders – in order to produce the resources necessary to fuel their collective success in intergroup conflicts, while freeing their leaders from carrying those prosocial norms over to their behavior in inter-group interactions. Less democratic groups are thereby able to leverage the resources generated by their members’ cooperative interactions by granting greater influence to those members who have accumulated the most resources – individuals who will inevitably be those with the least cooperative strategy traits. This mechanism facilitates the separation of the behavior of the collective from the behavioral norms of the majority of individuals populating the group.

In an interconnected world, the most successful groups will collectively take on the personality traits of their least cooperative members. Further to this point, we might begin to think of sub-categories of non-cooperating individuals – and personality traits characterized as Machiavellian, psychopathic, or narcissistic (the “dark triad”) – as minority traits selected for at the group level (Paulhus & Williams, 2002), lending a new twist to adaptationist theories of those traits (Wilson et al., 1996; Glenn et al., 2011; Gervais et al., 2013). Our model suggests that organizational frameworks that put more selfish individuals in charge of the group’s collective decisions may be favored, as least in so far as those decisions are related to cooperation and competition with other groups.

Although we predict that high interconnectedness will select for more plutocratic institutions of power, the inverse is not explicitly supported by our model. In cases of low interconnectedness, selection on group-level institutions of power is neutral, and therefore the ubiquity of democratic and egalitarian institutions among foraging societies must be explained through processes not fully captured by our model. Boehm (1999) has suggested that egalitarian institutions foster norms of cooperation and trust within the group, and serve to minimize fitness differences within the group while amplifying differences between groups, which facilitates group-level selection for cooperation in the manner described by multilevel selection theory (Wilson & Wilson, 2007). However, this cannot be the full story, as division of labor and the consequent inequality can also have group-level benefits (Henrich & Boyd, 2008; Smaldino, 2014). It is likely that the stationary lifestyles and wealth accumulation associated with the transition to agriculture at the dawn of the Holocene played some part in the transition from more egalitarian to more skewed institutions of power (Richerson & Boyd, 1999; Bowles & Choi, 2013; Gowdy & Krall, in press).

### 4.1. Limitations and future directions

Our model explores the cultural evolution of institutions of power in response to broad changes in between-group interactions. We do not provide a generative model for how these latter changes came about, nor do we make any specific claims about the specific state of the world either historically or presently. Indeed, our model applies just as readily to a world in which intergroup conflict is *decreasing.* Nevertheless, the historical and archaeological record supports a world in which human groups experienced dramatic increases in their rates of interaction with other ethnic and political groups (Johnson and Earle 2000; Horowitz 2001; Bowles 2009). These changes occurred via complex, multistage processes involving technological and infrastructural advances that are absent from our analysis. Because our model merely assumes increased rates of inter-group interaction without specifying mechanism, our results are entirely compatible with more specific mechanistic explanations for the evolution of social complexity

Modeling social evolution necessitates presenting a highly simplified picture of the world. We have not taken into account many important complexities, including norms of social enforcement, contingent or adaptive strategies of cooperation, or the roles of technology or social organization. Consideration of these factors underlies important questions that will form the basis of future work.

For simplicity, we considered only two levels of organization and interaction: the individual and group levels. While this is still one level more than most evolutionary population models include, real human social organization is quite a bit more complicated. Both large- and small-scale societies are hierarchically organized, with levels of organization ranging from individuals and families through bands to multi-band societies in the case of hunter-gatherers (Hamilton et al., 2007). These complexities in group structure will affect our characterization of power and how it evolves – the full story will involve the transition from simple patterns of asymmetry at the group level to power structures based on multiparty interactions and assessments to the emergence of formalized leadership positions (Boehm & Flack 2010). We also assume the existence of costless group markers. If group membership must instead be established through the use of costly signaling and sacrifices, such as those fostered by religious rituals (Iannaccone 1992; Aimone, Iannaccone et al. 2013; Makowsky 2012; Sosis, 2003), then the emergent social structures from those organizations may be opposed, both culturally and in terms of cost tradeoffs, to institutions of power that would be optimal in the absence of costly signaling.

Groups in our simulated world were initialized with a fixed institution of power. Details related to how these institutions evolve endogenously within groups or spread between groups were left out. Some models exist in which cooperative norms can spread through social learning in a group-structured population (e.g., Boyd & Richerson, 2002), and similar models may be useful for our purposes. A caveat, however, is that institutional evolution is complicated by the fact that the adoption of a new institution is a group-level process that requires widespread coordination. The details of warfare and resource allocation were also simplified for convenience. In reality, smaller groups will have access to fewer resources, the risk of group-level decisions may assessed before engaging in conflict, and allocation of the spoils of war or conquest among the individuals within a groups may be heavily weighted to those with more power. Future work will address details such as resource allocation, technology, and returns to scale. Finally, our model only considers collective decisions related to inter-group conflict, and does not address the many other group-level costs, benefits, and dynamics of various institutions of power (see Acemoglu, 2008; Makowsky and Rubin 2013; Rubin 2014).

### 4.2. Concluding remarks

Autocracy, plutocracy, aristocracy, and other variants of top-heavy distributions of power present a double-edged sword. Such skewed power distributions may facilitate an increasingly interconnected world through positive feedback, by freeing the majority of citizenry to be more cooperative in individual-level interactions, such as trade, with outsiders. While fostering
cosmopolitan cooperation norms, however, these social institutions co-evolve with conflict and aggression, imbuing the most selfish agents with the scale and power of their groups in contexts within which they can be the most potentially destructive.

## Supplementary Information

### S1. Algorithm for the initialization of group *powerGinis*

In order to generate a distribution of *powerGinis* that were nearly uniformly distributed between 0 and 1, we used the following algorithm to generate each group’s power skew parameter.

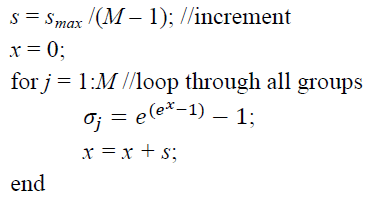

where *s*_*max*_ was a scale parameter set to 1.65. The resulting distribution of *powerGini* coefficients is illustrated in Fig. S1.

**Figure S1.**
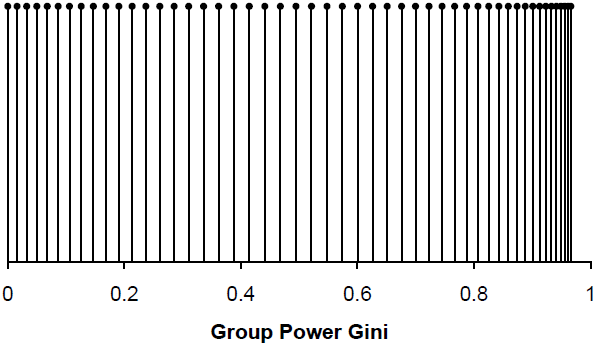
The initial distribution of group *powerGinis* for all groups.

### S2. A closer look at the emergent power structure

In the main text, we put forth the claim that inter-group conflict and skewed power structures facilitates individual-level cooperation by creating a leadership class of the selfish. Here we examine in more detail the differences in cooperative behavior between those individuals within a group who are the most or the least cooperative, and how unequal distributions of power facilitate the divergence of norms of group behavior (median and mean individual behavior) from leadership strategies (the least cooperative individuals in groups with higher *powerGini* coefficients).

Figure S2. and Figure S3 show aggregate data from across groups, depicting the relationship between rate of conflict, population mixing, and cooperation strategies. The results reaffirm that out-group cooperation generally decreases with the rate of population mixing, while in-group cooperation increases. It is notable that while out-group cooperation always increases with the rate of conflict, the correlation is much stronger for the most cooperative individuals (50^th^ percentile and greater), while the 1^st^ through 5^th^ percentiles remain largely unchanged (Fig. S2). Similarly, in-group cooperation always increases with the level of population mixing, and the correlation is much stronger for the most cooperative individuals (50^th^ percentile and greater), but the 1^st^ percentile again remains unchanged (Fig. S3). In tandem, population mixing and conflict serve to increase overall cooperation while at the same time widening the distribution of strategies.

**Figure S2.**
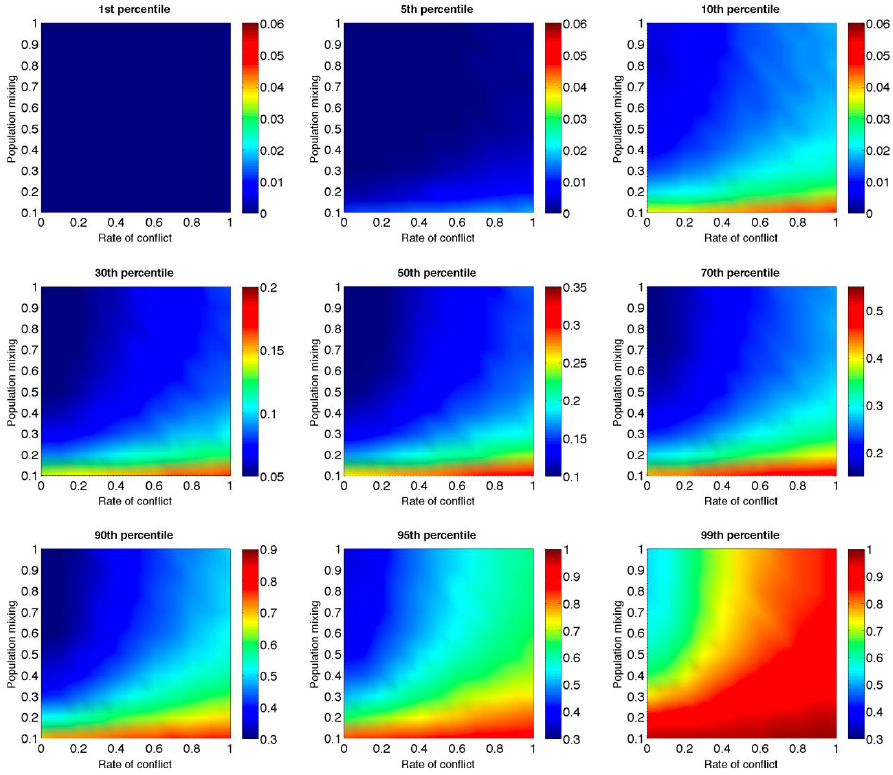
Out-group strategy distributions: the leadership of the selfish remains unchanged. Across groups, those individuals in the lowest percentile of out-group cooperation strategies remain unchanged by population mixing and conflict. The middle and upper portions of the distributions are affected the most. As noted earlier, conflict rate has a greater effect on out-group strategies than population mixing level.

**Figure S3.**
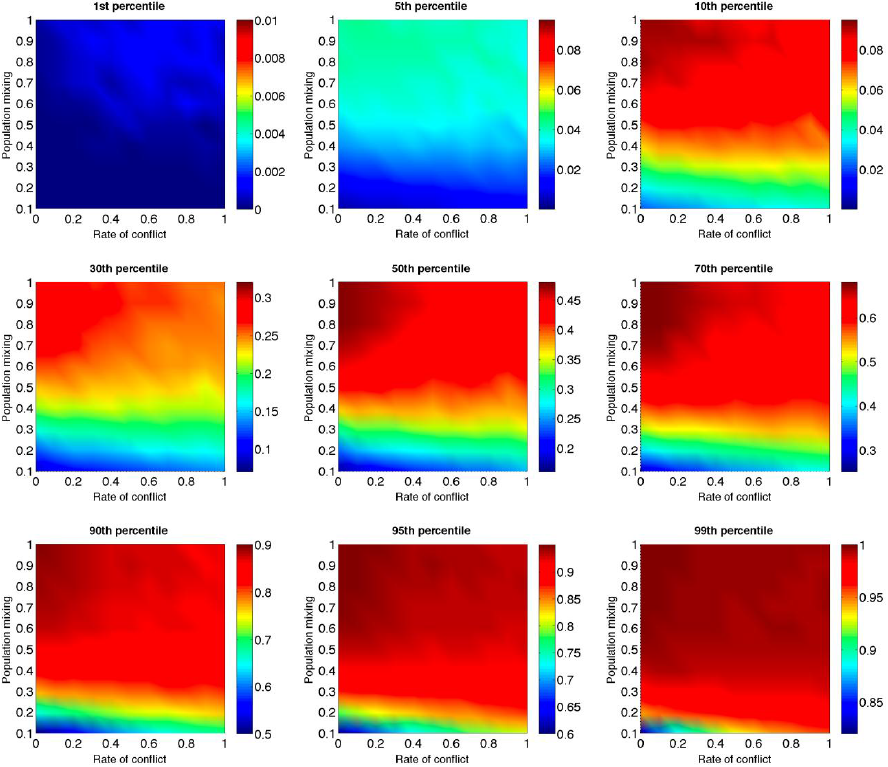
In-group strategy distributions: the leadership of the selfish remain unchanged. Across groups, those individuals in the lowest percentile of in-group cooperation strategies remain unchanged by population mixing and conflict. The middle of the distributions is affected the most. As noted earlier, the population mixing level has a greater effect on in-group strategies than conflict rate.

### S3. The absence of group markers

Humans cooperate and engage in collaborative ventures with a wide range of individuals, many of whom are not only non-kin, but may be complete strangers. One of the mechanisms through which we do so is by the maintenance of ethnic or group markers, which have been shown to increase both the capacity for cooperation (Sherratt & Roberts, 2001; Efferson et al., 2008; Fu et al., 2012) and for more complex collaborations involving coordination and distribution of labor (McElreath et al., 2003). Group markers may include physical attributes such as clothing, hairstyles, and makeup, as well as behavioral characteristics such as language and accent. Empirical evidence suggests that humans possess psychological machinery that facilitates group identification and the detection of coalitions and alliances (Kurzban et al., 2001); and that individuals often use these markers as a basis for choosing with whom to cooperate (Dunbar & Nettle 1997; Price 2006; Mathew & Boyd, 2011; Cohen & Haun 2013). Group markers are also crucial in the evolution of parochial altruism: you cannot preferentially cooperate with group members if you cannot tell them apart from outsiders (Choi & Bowles 2007; Garcia & van der Bergh 2011). We therefore assumed the existence of reliable group markers.

However, although the existence of group markers is not disputed, the applicability to cooperation is sometimes contested (Fehr & Fischbacher 2005; Gardner & West 2009). Low-cost group markers could be faked, rendering them useless. We do not believe this objection is particularly strong, given the widespread evidence on the use of group markers in human cooperation. Nevertheless, it is important that the basic results of our model – that inter-group conflict and skewed power structures facilitates individual-level cooperation by creating a leadership class of the selfish – was not dependent on group markers.

We ran simulations in the absence of group markers, so that individuals only had one genetic trait, *p*, which was the probability of cooperating on any interaction, regardless of whether one’s co-player belonged to one’s own group. Our results support previous findings that group markers facilitate increased cooperation, and in particular a dramatic increase in parochial altruism (Fig S4). Nevertheless, the qualitative results of inter-group conflict increasing cooperation still held, as did the effects seen in main text Fig 2B and 2C. The effect seen in main text Fig 3 also held in that higher rates of conflict led to greater success for groups with more autocratic power distributions, but, predictably, the rate of population mixing was no longer a factor in this equation.

**Fig S4.**
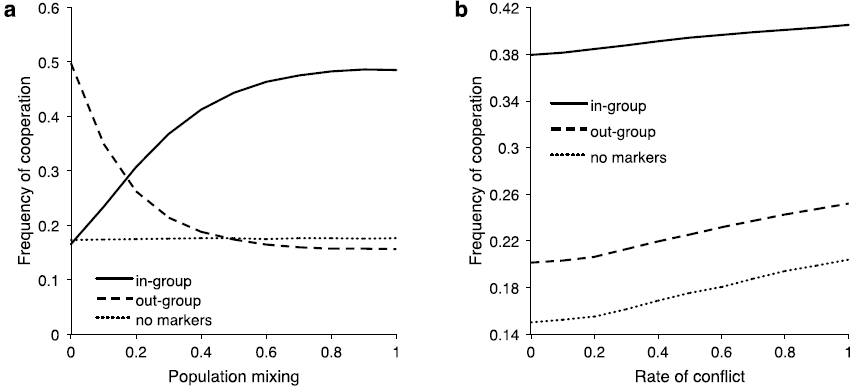
Group markers facilitate increased cooperation.

## Footnotes

1 This is not to say the individual interactions cannot be negative sum (e.g., theft) or that group interactions cannot be positive sum (e.g., trade agreements). Nevertheless, between-individual cooperation and inter-group conflict are widely considered to be among the most important factors, broadly speaking, in human social evolution (Choi and Bowles 2007; Hill, Barton et al. 2009; Bowles and Gintis 2011; Turchin, Currie et al. 2013; Richerson, Baldini et al. 2014).

2 Our theoretic model is built with individuals carrying a single strategy trait for out-group interactions, as opposed to separate strategies for personal and collective interactions. This is because the selection pressure on a collective behavior trait carried by an individual would have minimal fitness consequences for the individual. As such it would be (within a cultural evolution model) effectively randomized relative to individual behavior traits that are being selected for. This is likely our most controversial assumption. We would also note that this assumption minimizes the cognitive costs, and cognitive dissonance, relative to carrying two potentially opposing strategies, a non-trivial point given that boundedly rational agents will often generalize strategies learned in one context to another less familiar context (Simon 1956; Czerlinski, Gigerenzer et al. 1999; Bednar and Page 2007).

3 In models of cultural evolution, resource-based (i.e., payoff-based) reproduction is equivalent to success-biased imitation, in which the strategies of the most successful individuals are preferentially copied (Henrich and Gil-White 2001).

4 Another approach would be to initialize offspring with resources proportional to those of their parents. This assumption is perhaps more realistic. However, because offspring inherit the strategies of their parents, proportional resource allocation would create positive feedback in the evolutionary dynamics, giving much greater influence to the wealthiest individuals in autocratic societies. While this description may indeed reflect reality in at least some times and places, uniform allocation of resources to offspring provides a more rigorous test of our hypothesis, by requiring individuals to pull themselves up by their bootstraps, their pockets unburdened by their pedigree.

5 The model reaches a steady state – a state of broken ergodicity (Axtell 2000), if you will – well before step 300, typically between steps 150 and 200. While the relative distribution of population across groups reaches a steady state, the overall population continuously grows. 300 time steps was chosen as the cutoff because it was safely after a steady state was consistently reached, but before population growth reached a point of impractical computational cost. We should also note that while the difference in rates of cooperation between groups reach a steady state, all rates of cooperation across the entire population continue to decay, albeit at a slowing rate. This is largely a product of the simplicity of the model – there is no agent learning, monitoring, or punishment in the model, and no mechanism for purposely repeated interactions.

6 It is important to note that migration, in this context, is simply a form of noise added to the model to insure against knife-edge results. A more meaningful model of migration would require purposeful movement and destination decisions on the part of agents.

7 Within groups, selfish agents will outcompete cooperative agents only up to the point where their selfishness might cause the entire group to go extinct, which contributes to the persistence of cooperative strategies in our model. Moreover, the presence of iterated (selectively repeated) games would change the evolutionary dynamics, because pairs of cooperators could outperform defectors. For simplicity, we have not explored this additional layer of model complexity.

